# Speciation in a biodiversity hotspot: phylogenetic relationships, species delimitation, and divergence times of the Patagonian ground frogs of *Eupsophus roseus* group (Alsodidae)

**DOI:** 10.1101/423202

**Authors:** Elkin Y. Suárez-Villota, Camila A. Quercia, Leila M. Díaz, Victoria A. Vera-Sovier, José J. Nuñez

## Abstract

The alsodid ground frogs genus *Eupsophus* is divided into the *roseus* (2n=30) and *vertebralis* (2n=28) groups, distributed throughout the temperate *Nothofagus* forests of South America. Currently, the *roseus* group is composed by four species, while the *vertebralis* group consists of two. Phylogenetic relationships and species delimitation within each group are controversial. In fact, previous analyses considered that *roseus* group was composed between four to nine species. In this work, we evaluated phylogenetic relationships, diversification times, and species delimitation within *roseus* group using a multi-locus dataset. For this purpose, mitochondrial (D-loop, Cyt *b*, and COI) and nuclear (POMC and CRYBA1) partial sequences, were amplified from 164 individuals, representing all species. Maximum Likelihood (ML) and Bayesian approaches were used to reconstruct phylogenetic relationships. Species tree was estimated using BEAST and singular value decomposition scores for species quartets (SVDquartets). Species limits were evaluated with six coalescent approaches. Diversification times were estimated using mitochondrial and nuclear rates with LogNormal relaxed clock in BEAST. Nine well-supported monophyletic lineages were recovered in Bayesian, ML, and SVDquartets, including eight named species and a lineage composed by specimens from Villarrica population (Bootstrap: >90, PP:> 0.9). Single-locus species delimitation analyses overestimated the species number in *E. migueli, E. calcaratus* and *E. roseus* lineages, while multi-locus analyses recovered as species the nine lineages observed in phylogenetic analyses (>0.95). It is hypothesized that *Eupsophus* diversification occurred during Mid-Pleistocene (0.42-0.14 Mya), with most species originated after of the Last Southern Patagonian Glaciation (0.18 Mya). Our results revitalize the hypothesis that *E. roseus* group is composed by eight species and support to Villarrica lineage as a new putative species.

## Introduction

From the operational point of view, the notion of biodiversity encompasses several different levels of biological organization, from the species’ make up genetic to ecosystems and landscapes, in which the species is the most significant unit. Species are used for comparisons in almost all biological fields including ecology, evolution, and conservation [1–3]; no doubt the central unit for systematics is also the species [4]. Furthermore, biodiversity hotspots are selected on the basis of the species they possess, conservation schemes are assessed on how many species are preserved, and conservation legislation and politics are focused on species preservation [5,6].

Although the importance of species concepts debate [7,8], and that the species as taxonomic hierarchy is also considered a fundamental topic in biology [9], it is broadly accepted that species are best conceptualized as dynamic entities, connected by “grey zones” where their delimitation will remain inherently ambiguous [4,10]. Under this perspective, species delimitation, i.e. the act of identifying species-level biological diversity [11], is particularly challenging in actively radiating groups composed of recently diverged lineages. The difficulty lies in the fact that recently separated species are less likely to possess all or even many of the diagnosable characters such as phenetic distinctiveness, intrinsic reproductive incompatibility, ecological uniqueness, or reciprocal monophyly, that constitute operational criteria for their delimitation [4,12]. Thus, hypotheses of the boundaries of recently diverged species can remain unclear due to incomplete lineage sorting, introgression, complex of cryptic species that cannot be distinguished by morphology alone, sampling deficiencies, or different taxonomic practices [2,4].

As genetic data have become easier and less expensive to gather, the field of species delimitation has experienced an explosion in the number and variety of methodological approaches [3,11,13–15]. These new approaches proceed by evaluating models of lineage composition under a phylogenetic framework that implements a coalescent model to delimit species [11,16]. In this regard, these approaches estimate the phylogeny while allowing for the action of population-level processes, such as genetic drift in combination with migration, expansion, population divergence, or combinations of these processes [17–19]. Thus, species delimitation models can involve population size parameters (i.e. θs for the extant species and common ancestors), parameters for the divergence times (τ), and coalescent models specifying the distribution of gene trees at different loci [20–24].

Some methodological approaches to species delimitation use single-locus sequence information itself as the primary information source for establishing group membership and defining species boundaries [25–27]. Other methods are designed to analyze multi-locus data sets and require a priori assignment of individuals to species categories [19,28,29]. The performance of species delimitation methods are quantified by the number of different species recognized in each case and the congruence with data at hand as life history, geographical distribution, morphology, and behavior [13,30]. Although, there is difficulty to integrate genetic and non-genetic data to increase the efficacy of species detection [31], there are available methods to measure the congruence and resolving power among species delimitation approaches [32].

Patagonian landscape history offers exceptional opportunities to investigate diversification and promote conservation strategies by studying past, present, and future of evolutionary processes using amphibians as model study. In this region, the amphibian fauna of Chile is not particularly diverse (60 species; [33]), but includes 10 endemic genera, some of them having one or few species (e. g. *Calyptocephalella, Chaltenobatrachus, Hylorina, Insuetophrynus, Rhinoderma*), to as many as 18 (*Alsodes*). Among these amphibians are frogs of the genus *Eupsophus* Fitzinger 1843. This taxon includes currently six species distributed almost throughout the temperate *Nothofagus* forest of South America [33]. Nevertheless, *Eupsophus* have puzzled frog systematics for decades [34–37], and a clear consensus has not yet been reached regarding the number of species that make up this genus [38–40]. In fact, the genus *Eupsophus* was classically divided into two groups with following species [34,41]: 1) *roseus* group, composed of *E. altor, E. roseus, E. calcaratus, E. contulmoensis, E. insularis, E. septentrionalis, E. migueli* and *E. nahuelbutensis*. All of them with 30 chromosomes, and whose individuals have a body size of 34-42 mm (snout-vent distance) [42]; and 2) the *vertebralis* group, composed of *E. vertebralis* and *E. emiliopugini*, both species with 28 chromosomes and individuals with a body size of 50-59 mm (snout-vent distance) [42]. Nevertheless, recently molecular analyses within *roseus* group synonymized *E. altor* with *E. migueli* as well as *E. contulmoensis, E. septentrionalis*, and *E. nahuelbutensis* with *E. roseus* [35]. Therefore, currently the *roseus* group is composed by four species: *E. migueli, E. insularis, E. roseus* and *E. calcaratus* [33].

Here, we present phylogenetic and species delimitation of the *roseus* group, using 164 new samples from all species covering most of their distribution range. We used three mitochondrial and two nuclear markers, three of them are different to those used by Blotto et al [34] and Correa et al [35] [Control Region (D-loop), Propiomelanocortin (POMC), and β Crystallin A1 (CRYBA1)]. These molecular dataset are used to carry out phylogenetic reconstructions and an extensive number of single- and multi-locus species delimitation methods. Species trees and diversification times were estimate to support phylogenetic and species boundaries inferences. New samples, different markers, and multiple bioinformatic techniques allowed us to test, in an independent way, phylogenetic and species delimitation hypothesis in the *roseus* group.

## Materials and Methods

### Ethics Statement

This study was carried out in strict accordance with the recommendations of the Bioethics and Biosecurity Committee of the Universidad Austral de Chile (UACh), Servicio Agrícola y Ganadero (Resolución ExentaN° 9244/2015). After capture, animals were kept in the dark in fabric bags for a maximum of two hours. Euthanasia was carried out in the field via intra-abdominal injection of sodium pentobarbital at a dosage of 100 mg/kg of body weight. The Corporación Nacional Forestal, Ministerio de Agricultura, Gobierno de Chile allows to collect buccal swabs samples of *Eupsophus* species from wild protected areas (Permit No. 11/2016.-CPP/ MDM/jcr/ 29.02.2016).

### Sample collection

In total, 164 samples of *Eupsophus* from 45 localities in Chile were analysed (Fig 1, S1 Table). Each sampling site was geo-referenced with a GPS Garmin GPSmap 76CSx. Two individuals of *E. emiliopugini*, three *E. vertebralis*, and one *Alsodes valdiviensis* were used as outgroup (S1 Table, gray cells). Although mostly samples were obtained from buccal swabs according to Broquet et al. [43], some animals were euthanized. Liver tissue was extracted, conserved in 100% ethanol, and stored at −20°C. Specimens were deposited in herpetological collection from Instituto de Ciencias Marinas y Limnológicas, Universidad Austral de Chile (ICMLH). Voucher and isolate numbers were included in sequences information.

**Fig 1.**
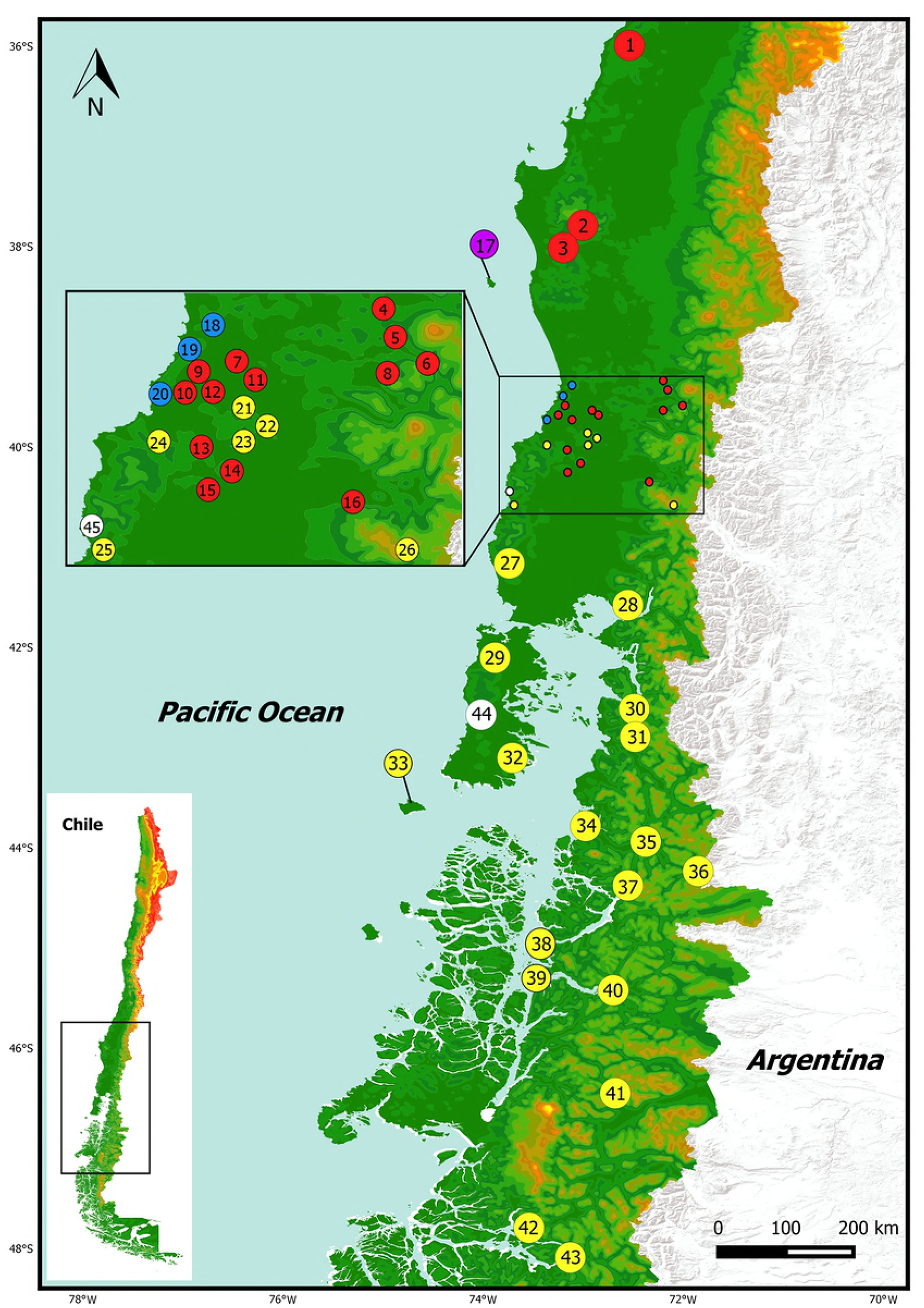
Map depicting 45 localities of *Eupsophus* samples from Chile (listed in S1 Table). *E. roseus*: localities 1-16 (red), *E insularis*: locality 17 (purple), *E. migueli*: localities 18 - 20 (blue), *E. calcaratus*: localities 21-43 (yellow). Localities of outgroup were: *E. emiliopugini*: 44 and 45 (white), *E. vertebralis*: 12, 19, 22, *Alsodes norae*: 19.

### DNA extraction, amplification, and sequence alignment

Whole genomic DNA was extracted using Chelex following Walsh et al. [44]. We amplified via the polymerase chain reaction (PCR) three mitochondrial regions: a segment of D-loop [45], Cytochrome *b* (Cyt *b*; [46]), and Cytochrome oxidase subunit I (COI; [47]), and two nuclear regions: POMC [48], and CRYBA1 [49]. We mixed reaction cocktails for PCR using 100 ng DNA, 10 µmol of each oligonucleotide primer, 2X of Platinum^®^ *Taq* DNA Polymerase master mix (Invitrogen, Cat. No. 10966), and nuclease-free water to final volume of 25 μL. We verified successful PCR qualitatively by viewing bands of appropriate size following electrophoresis on 1.0% agarose gels. PCR products were sequenced in Macrogen Inc. (Seoul, Korea). Electropherograms were visualized and aligned with Geneious v.9.1.3 (GeneMatters Corp.) using the iterative method of global pairwise alignment (Muscle and ClustalW) implemented in the same software [50,51]. An inspection of aligned sequences by eye and manual corrections were also carried out. All sequences from *Eupsophus* and *Alsodes* were submitted to Genbank (XX000000-XX00000).

### Phylogenetic analyses

Phylogenetic trees were constructed with concatenated dataset using Maximum Likelihood (ML) and Bayesian inference (BI). Evolutionary models and partitioning strategies were evaluated with Partitionfinder v2.1.1 [52] and the best partition was identified using the Bayesian information criterion [53]. ML trees were inferred using GARLI v2.0 [54] with branch support estimated by nonparametric bootstrap (200 replicates) [55]. Bayesian analyses were performed using MrBayes v3.2 [56]. Each Markov chain was started from a random tree and run for 5.0×10^7^ generations with every 1000th generation sampled from the chain. MCMC stationarity was checked as suggested in Nylander et al. [57]. All sample points prior to reaching the plateau phase were discarded as “burn-in”, and the remaining trees combined to find the a posteriori probability of phylogeny. Analyses were repeated four times to confirm that they all converged on the same results [58].

Species tree were reconstructed using the Singular Value Decomposition Scores for Species Quartets (SVDquartets) [62] and species tree reconstruction in BEAST v2.4.8 (*BEAST) [28,63].

SVDquartets method infers relationships among quartets of taxa under a coalescent model and then estimates the species tree using a quartet assembly method [59,60]. We evaluated all the possible quartets from the concatenated data set using SVDquartets module implemented in PAUP* v4.0a [61]. Quartet’s Fiduccia and Mattheyses algorithm [62] and multispecies coalescent options were used to infer species tree from the quartets. We used nonparametric bootstrap with 100 replicates to assess the variability in the estimated tree [55].

For *BEAST, multi-species coalescent module implemented in BEAST [28,63] and concatenated dataset were used. We set the partition scheme and models found by Partitionfinder. Mutation rates, clock models, and tree priors were the same detailed in divergence time estimates section (see below). MCMC were run three times for 5.0×10^7^ generations each, logging tree parameters every 50,000 generations. Posterior distribution was summarized with Densitree v2.01 [63]. Chain mixing, convergence, and a posteriori probability were estimated in the same way of the Bayesian analyses described above.

### Species delimitation analyses

Two single-locus analyses, Bayesian General Mixed Yule Coalescent model (bGMYC; [25,64]) and multi-rate Poisson Tree Processes (mPTP; [65]) were performed on mitochondrial dataset. The GMYC model distinguishes between intraspecific (coalescent process) and interspecific (Yule process) branching events on a phylogenetic tree [27]. We used the last 100 trees sampled from the posterior distribution of a Bayesian analysis for mitochondrial sequences (detailed in next section). Bayesian GMYC analyses were assessed using the R package bGMYC, where each tree was ran for 50,000 generations, discarding the first 40,000 generations as burn-in and using thinning intervals of 100 generations (as recommended by Reid and Carstens [66]). The threshold parameter priors (t1 and t2) were set at 2 and 170, and the starting parameter value was set at 25.

mPTP is a phylogeny-aware approach that delimits species assuming a constant speciation rate with different intraspecific coalescent rates [65]. For this analysis, a tree obtained with mitochondrial dataset in MrBayes was used as input on the web server (http://mptp.h-its.org/#/tree).

Four multi-locus coalescent-based methods were applied to species delimitation: Tree Estimation using Maximum likelihood, (STEM; [16,19]), Bayesian Species Delimitation (BPP; [24,67]), Multi-locus Species Delimitation using a Trinomial Distribution Model (Tr2; [68]), and Bayes factor delimitation (BFD; [69]). As required by these software, a set of analyses assigning individuals to a series of species categories were performed (delimitation scenarios).

STEM analysis followed Harrington and Near [29]. ML scores for each species tree were generated with STEM v2.0 [19] and evaluated using information-theoretic approach outlined by [16].

BPP analysis was applied using Bayesian Phylogenetics and Phylogeography software (BPP v.2.2; [23,70]). We used A10 mode, which delimits species using a user-specified guide tree (species delimitation = 1, species tree = 0). Species tree obtained with *BEAST was used as guide tree. Population size parameters (θs) and divergence time at the root of the species tree (τ0) were estimated using A00 mode [67], while the other divergence time parameters were considered as the Dirichlet prior ([24]: equation 2). Each analysis was run four times to confirm consistency among runs. Following a conservative approach, only speciation events supported by probabilities larger or equal to 0.99 were considered for species delimitation.

Tr2 analysis followed Fujisawa et al. [68]. Gene trees were obtained in GARLI and its polytomies were resolved using internode branch lengths of 1.0×10^-8^ in Mesquite v2.75 [71].

For BFD analysis, we reconstructed a species tree for each delimitation scenario using BEAST, as it was detailed in phylogenetic analyses section (see above). After the standard MCMC chain has finished, marginal likelihood estimation (MLE) was performed for each species tree, using both path sampling and stepping-stone via an additional run of ten million generations of 100 path-steps (1,000 million generations). Subsequently, Bayes factor between delimitation scenarios were calculated using MLEs [69] and evaluated using the framework of Kass and Raftery [72].

The taxonomic index of congruence (*Ctax*) between pairs of species delimitation methods was estimated following Miralles and Vences’ protocol [32]. In order to access most congruent species delimitation approaches, mean *Ctax* value for each method was also estimated.

### Divergence time estimates

Divergence times were estimated with concatenated mitochondrial and nuclear dataset using the Bayesian method (BEAST v2.4.8; [63]). We used Neobatrachian mutation rates of 0.291037% and 0.374114% per million years for COI and POMC, respectively [73]. Mutation rates from the other markers were estimated using as prior nuclear or mitochondrial rates for all genes reported by Irrisarri et al. [73] (0.379173% and 0.075283% respectively). Partitionfinder provided nucleotide substitution models. LogNormal relaxed clock model and birth-death process as tree prior were used. Bayes factor analysis [74] indicated that this setting received decisive support compared with other models and tree priors availables in BEAST. Markov chains in BEAST were initialized from the tree obtained from species tree analyses to calculate posterior parameter distributions, including the tree topology and divergence times. We run this analyses for 5×10^7^ generations, and sampling every 1000th generation. The first 10% of samples were discarded as “burn-in”, and we estimated convergence to the stationary distribution and acceptable mixing using Tracer v1.6 [75]. An additional BEAST analysis was carried out with only mitochondrial dataset using the same setting to obtain the last 100 trees. These trees were used as input in bGMYC (see section above).

## Results

### Phylogenetic patterns in E. roseus group

We aligned the five DNA markers for a total of 2576 sites, 858 were variable and 700 were phylogenetically informative. Three of these markers corresponded to mitochondrial dataset with a total of 1799 nucleotide sites, 750 variable, and 629 phylogenetically informative (see information for each marker in S2 Table). Evolutionary models and partitioning strategy obtained in Partitionfinder are also indicated in supplementary data (S2 Table).

The phylogenetic analysis using concatenated mitochondrial and nuclear sequences recovered three main well-supported clades corresponding to Clade A (including *E. insularis* and *E. migueli*), Clade B (*E. roseus*) and Clade C (*E. calcaratus*) (Fig 2). Although ML and Bayesian analyses recovered to B and C were sister clades, phylogenetic relationships among these clades received low support (Fig 2). Within these clades is possible recognize nine highly supported monophyletic lineages (Fig 2; Bootstrap >90, PP>0.9, lineages 1-9).

**Fig 2.**
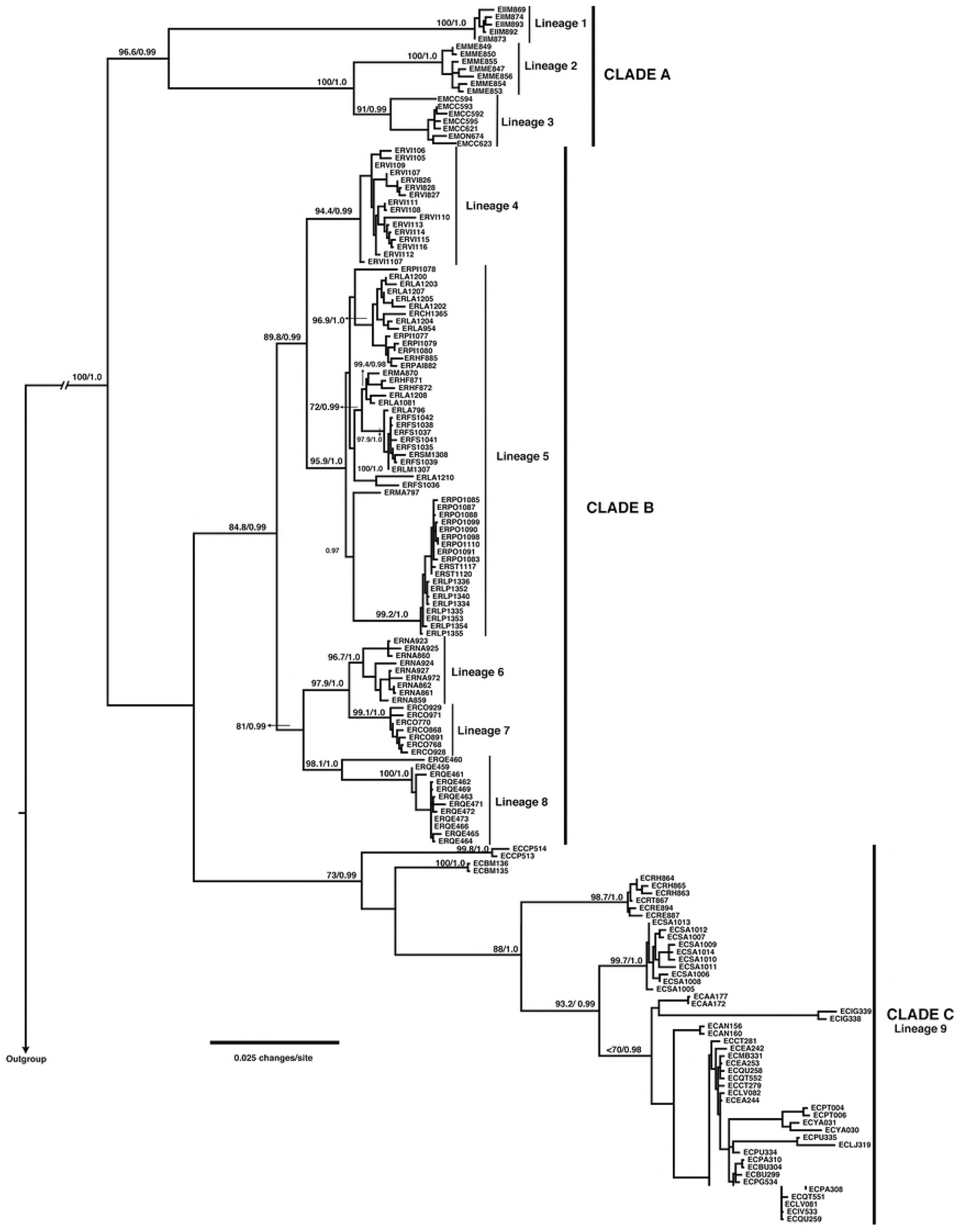
Phylogenetic relationships among *Eupsophus* species. This maximum likelihood (ML) tree was reconstructed using concatenated nuclear and mitochondrial data set. Topologies obtained by ML and Bayesian inference were similar. Numbers above branches represent bootstrap scores and Bayesian posterior probabilities. Isolate numbers consist by the species abbreviation (*E. roseus*: ER, *E. migueli*: EM, *E. insularis*: EI, and *E. calcaratus*: EC), locality abbreviation listed in S1 Table, and field number. Major clades (A, B, and C) and lineages (1-9) of *Eupsophus* are indicated.

### Species delimitation analyses

The most congruent result among single- and multi-locus analyses recognized nine monophyletic lineages as different species (Fig 3; mean *Ctax*= 0.69, see all *Ctax* values in S3 table). These nine lineages were the same recovered in the phylogenetic analyses and were also supported in the consensus tree from the SVDquartets analysis (Fig 3; Bootstrap >70). Having in mind, geographical distribution (Fig 1) and phylogenetic analyses of Blotto et al [34], these lineages corresponded to the formerly eight *Eupsophus* species of the *roseus* group: *E. altor, E. migueli, E. insularis, E. contulmoensis, E. nahuelbutensis, E. septentrionalis, E. roseus, E calcaratus*, plus a lineage composed by specimens from Villarrica locality, hereafter referred to as *Eupsophus* sp. (Fig 3).

**Fig 3.**
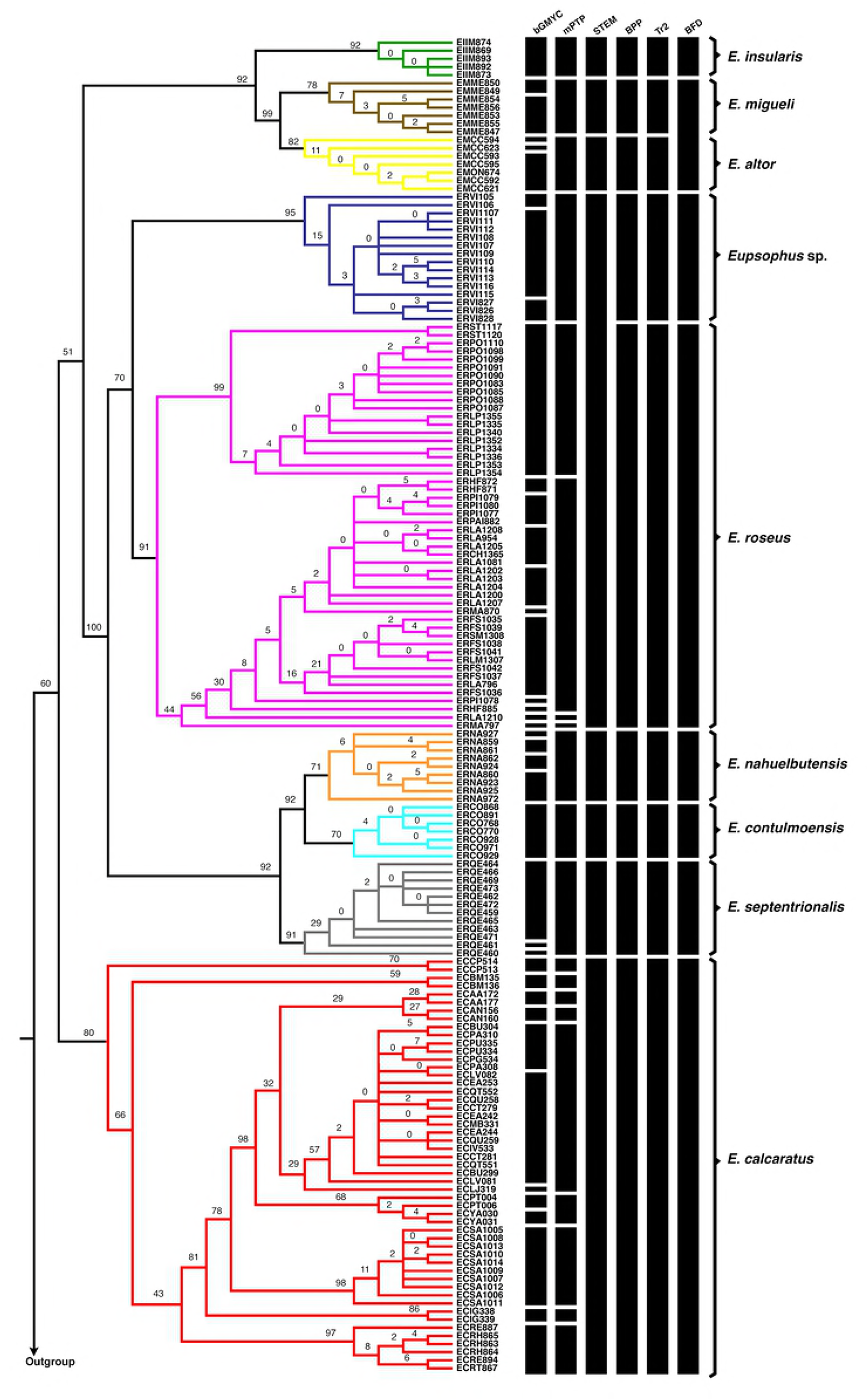
SVDquartets and species delimitation analyses. Majority-rule consensus tree from the SVDquartets analysis. Nodal support values are bootstrap proportions. Bars on the right of the tree indicate the species limits as proposed by bGMYC, mPTP, STEM, BPP, Tr2 and BFD analyses. All analyses were carried out with mitochondrial and nuclear loci, except bGMYC and mPTP which used only mitochondrial data set. Limits of formerly *Eupsophus* species and putative species from Villarrica (*Eupsophus* sp.) are indicated with different colors on the branches of the tree and with square bracket on the right of the bars. This limits correspond to the most congruent species delimitation scenario (see S2 table)

Bayesian GMYC analyses detected more than one species in these nine lineages except in *E. insularis* and *E. contulmoensis* (Fig 3). Multi rate PTP detected six species corresponding to *E. altor, E. migueli, E. insularis, E. contulmoensis, E. nahuelbutensis, E. septentrionalis* lineages, and more than one species in *E. roseus* and *E calcaratus* lineages (Fig 3). Nine-species scenario (Fig 4A, gray cell) was the highest supported in BPP and Tr2 analyses (Fig 4B, black arrows, scenario 12). For STEM analysis the eight-species scenario, where *Eupsophus* sp. and *E. roseus* represent a single species, was the highest supported (Fig 4A, scenario 11). Nevertheless, among the other species delimitation scenarios, the STEM analysis greatly favored a nine-species delimitation scenario (Fig 4B, S4 Table). Highest MLEs in BFD analysis were obtained for eight-species scenario, where *E. altor* and *E. migueli* corresponded to one species (Fig 4, scenario 10). In this case, Bayes factor comparisons were greater than two, which allowed us to choose that better scenario (S5 Table). Nevertheless, comparisons with some scenarios including that of nine-species were around four, which indicate non-strong or decisive support to the best model (S5 Table). Other possible scenarios, including that proposed by Correa et al. [35] (scenario 3), were lowly supported for all multi-locus analyses (Fig 4).

**Fig 4.**
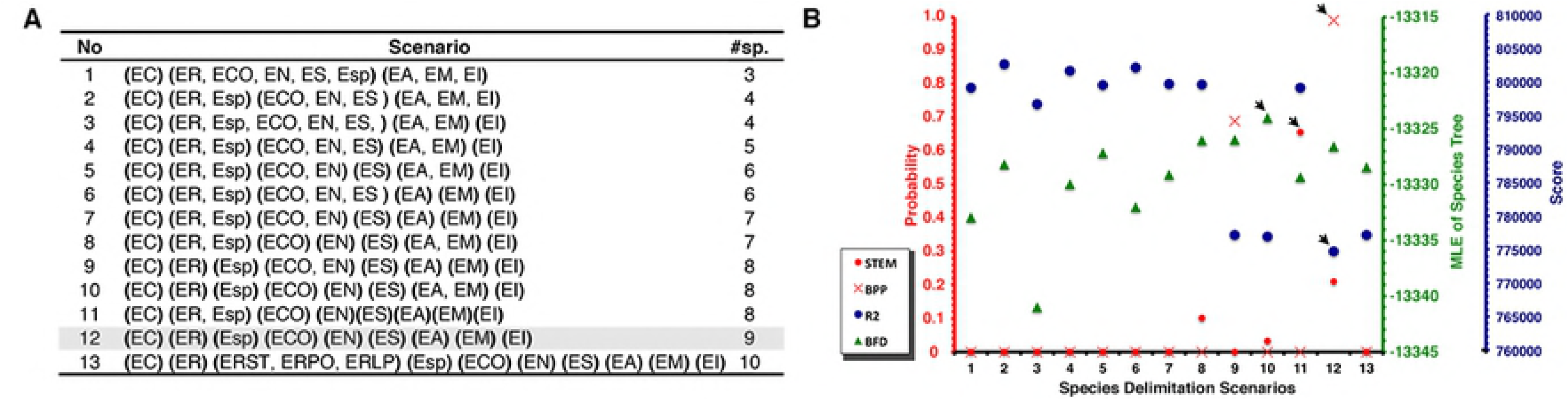
Multi-locus species delimitation analyses. A) species delimitation scenarios. Specimens were assigned to delimited species indicated in Fig 3. Abbreviations within parenthesis indicate the grouping tested in each scenario. *E. roseus*: ER, *E. migueli*: EM, *E. insularis*: EI, and *E. calcaratus*: EC, *E. altor*: EA, *E. contulmo*: ECO, *Eupsophus* sp.: EV, *E. nahuelbutensis*: EN, *E. septentrionalis*: ES. Some abbreviated localities from S1 Table were added to species abbreviation to indicate a specific locality grouping. Most congruent scenario is indicated in gray. **B)** probability, marginal likelihood (MLE), or score values generated for each scenario using different species delimitation approaches. Black arrow indicates the credible species hypotheses. For Tr2 lowest score indicates the better-delimited scenario. For STEM and BFD were plotted model probabilities and MLE values using stepping-stone sampling, respectively (see S4 and S5 Tables)

### Species tree and divergence times estimates among Eupsophus species

Species tree reconstructions in *BEAST and SVDquartets, using the nine lineages (=species), recovered similar phylogenetic relationships to the Bayesian and ML analyses (Fig 5). Under this scenario, *E. calcaratus* diverged early in *Eupsophus* radiation for both species tree and divergence time tree. This topology appeared to be supported as it is revealed by overlaying posterior sets of trees generated by BEAST and plotted by DensiTree (Fig 5). Thus, we decided to used consensus species tree as a prior to estimate divergence times among *Eupsophus* species (Fig. 5, in blue).

**Fig 5.**
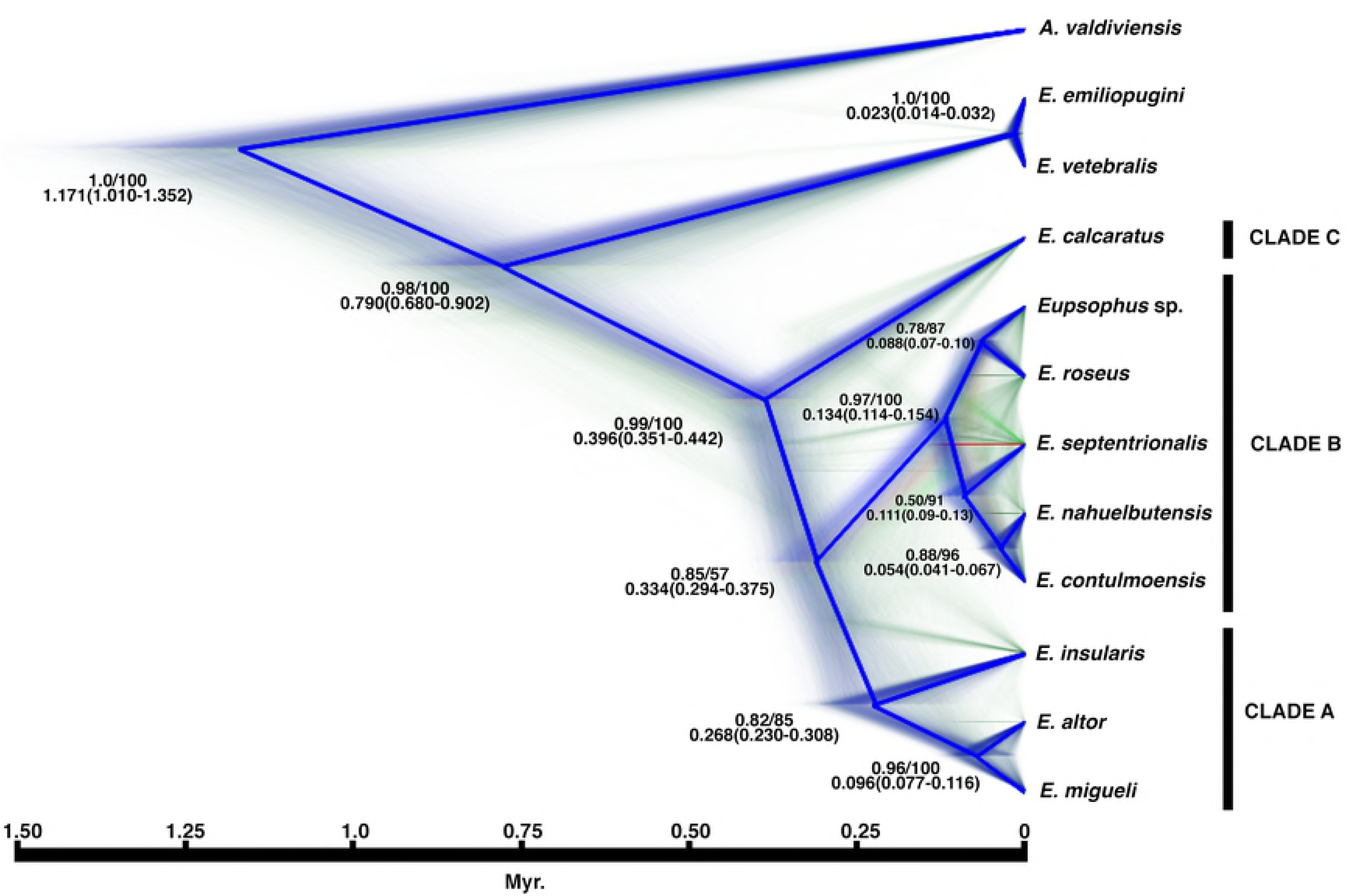
Species tree and divergence times of *Eupsophus*. This cladogram illustrates the posterior distribution of species trees inferred with BEAST based on the most congruent species delimitation scenario (Figs 3 and 4, S2 Table). High colour density is indicative of areas in the species trees with high topology agreement. Different colours represent different topologies. Consensus species tree are coloured in blue. Nodal values are Bayesian posterior probability (BEAST) and bootstrap proportions (SVDquartets). Mean divergence dates in million years and 95% credible intervals are indicated (below the support values).

The age of crown-group *Eupsophus* and the origin of *E. calcaratus* are estimated at 0.396 (0.351–0.442) Myr. *Eupsophus insularis* diverged at 0.268 (0.230–0308) Myr, while *E. altor* and *E. migueli* at 0.096 (0.077–0.116) Myr (Fig 5). The split between *E. roseus* and *Eupsophus* sp. /*E. contulmoensis, E. nahuelbutensis*, and *E. septentrionalis* was around 0.134 (0.114–0.154) Myr. The divergence between *E. roseus* and *Eupsophus* sp. is estimated at 0.088 (0.072–0.106) Myr. *Eupsophus septentrionalis* diverged at 0.111 (0.193–0.131) Myr, followed of *E. contulmoensis* and *E. nahuelbutensis* at 0.054 (0.041– 0.067) Myr (Fig 5).

## Discussion

### Species delimitation in the Eupsophus roseus group

The most congruent species delimitation results detected nine species in the *E. roseus* group, eight of them (namely *E. altor, E. calcaratus, E. contulmoensis, E. insularis, E. migueli, E. nahuelbutensis, E. roseus*, and *E. septentrionalis*), concordant with taxonomic proposals of the last decades [34,36,76–81].

The highest level of congruence was obtained with BPP and Tr2 methods (mean C*tax*=0.69; nine species), followed by STEM, and BFD (mean Ctax=0.63; eight species; Figs 3 and 4, S3 Table). Although, *Eupsophus* sp. and *E. roseus* clades were recovered as a single species by STEM, these clades were recovered as different species by BPP, Tr2, mPTP and BFD analyses. Similarly with *E. migueli* and *E. altor*, which were recovered as a single species by BFD but as two different species in the other analyses. Therefore, the greatest congruence indicate Clade B is composed by five different species (*Eupsophus* sp., *E. roseus, E. nahuelbutensis, E. contulmoensis* and *E. septentrionalis*), while Clade A by three (*E. altor, E. migueli,* and *E. insularis*) as it is suggested in previous works [78,82]. The differences among results of these species delimitation methods could be derived from its different sensibility to the ratio of population size to divergence time, such as it has been reported between BPP and bPTP [15]. Hence the importance of carrying out several species delimitation methods to examine whether the proposed groups are consistently recovered with different algorithms [15,11]. This is evident when we compared results from multi-locus analyses with bGMYC result (mean Ctax=0.27), which overestimated the species number in all lineages except in *E. insularis* and *E. contulmoensis* (Fig 3). It is known that bGMYC has shortcomings when datasets consist of few putative species [83] and cannot be used as sufficient evidence for evaluating the specific status without additional data or analyses [84]. Moreover, this method tends to overestimate the number of species when the ancestral polymorphism is low [85]. Therefore, rather than use this method as a species delimitation approach, we used it to obtain alternative scenarios to be tested with multi-locus analyses (e.g. scenario 13, Fig 4).

Our delimitation results were not agreed with a recent hypothesis [35], which would be related to use of different molecular markers and species delimitation analyses. Three of our markers were found to be highly variables (Cyt *b*, COI, D-loop), while two were conserved (POMC and CRYBA1; see S2 table). Thus we use at least three strong markers (sequences with many polymorphic sites), a key aspect to carried out coalescent analyses when less than ten markers are used [86]. On the other hand, we used several multi-locus coalescent methods to delimitate species (BPP, STEM, R2, and BFD), while Correa et al. [35] based its inferences in single-locus analyses (GMYC, mPTP, and Automatic Barcode Gap Discovery, ABGD). In this sense, mPTP (using mitochondrial data set) and ABGD (using mitochondrial + nuclear data set) recovered to the two groups of sinonimized species as two species [35]. ABGD method is based on genetic distances computed from a single-locus (COI) and requires a priori specification of an intraspecific distance threshold [87]. The robustness and accuracy of coalescent approaches over distance methods is well know, partly because the last do not appeal to an explicit species concept [15,88]. Therefore, we decided not to include ABGD in our main species delimitation analyses. Nevertheless, we conducted ABGD analyses using our COI data set, and our concatenated data set, obtaining different results (see S1 File). On this regard, using two potential barcode gaps, we detected nine and five groups with COI, while six and four groups were obtained with concatenated dataset. Consequently, ABGD results can be influenced by the application of a method designed for single-locus (DNA barcoding) to concatenated dataset, as well as by the a priori election of distance threshold. Moreover, ABGD analysis underestimated species diversity among species with low divergence [87,89]. Thus, ABGD tool is recommended as a first grouping hypothesis but not as robust and definitive species delimitation proof [87].

### Phylogenetic relationships and divergence time in the Eupsophus roseus group

Monophyly of *E. roseus* group and its nine delimited species was strongly supported, concordant with previous analyses (Fig 2; [34,82]. Although the early divergence of *E. calcaratus* was not strongly supported in Bayesian, ML, and SVDquartet approaches, our analyses resolved all other interspecific relationships among delimited species (Figs 2 and 3). In fact, the plot of overlying posterior sets of species trees (Fig 5) showed few alternative interspecific relationships. One example of this, is the early divergence of *E. septentrionalis* within Clade B, which was also recovered by Blotto et al [34] and Suárez-Vilota et al [81] (Fig 5, in red).

Phylogenetic and species delimitation analyses recognized to *Eupsophus* sp. as a distinct species (Figs 3 and 4). In fact, SVDquartet analysis detected this clade with greater support than other well-defined species such as *E. insularis* (Fig 3; bootstrap: 95%), and high probabilities were detected in single- and multi-locus species delimitation analyses (Fig 3 and 4). These results are concordant with previous works where suggested a species-level for this lineage [82]. Although Correa et al. [35] also detected a close phylogenetic relationship between Villarrica and *E. roseus* specimens, they considered the three specimens from this locality within the *E. roseus* diversity. We sampled 17 specimens from this locality and they were monophyletic with high support (Fig 2; Bootstrap: 100, PP: 1.0). Additionally, we did not detect syntopy instances in Villarrica, which could result in to recover specimens from other localities within Villarrica clade (i.e. interpopulational paraphyly). This paraphyletic pattern is common for localities within *E. roseus* lineage, an additional support to consider that Villarrica specimens do not belong to *E. roseus* species. For example, specimens from Fundo Santa María (FS) are recovered with specimens from other localities [e.g. Mafil (MA), Llancahue (LA)], in several highly supported clades within *E. roseus* lineage (Fig 2).

Mostly of delimited species from *E. roseus* group diverged from 0.134 to 0.054 Mya during Valdivian interglacial [90], except *E. calcaratus* and *E. insularis,* whose origin is older (before of the last southern Patagonian glaciation, 0.18 Mya). The oldest deposits of Mocha Island are dated from the Eocene and Miocene [91] whereas extensive terraces from Pliocene and Pleistocene characterize more recent settings [92]. Although the origin and presence of *E. insularis* in the Mocha Island remains unknown, these large terraces might have been a suitable habitat for its settlement and for its differentiation from the continental *Eupsophus* species. Anyway, it is possible that all species lived during Valdivia interglacial and subsequently were affected by the Last Glacial Maximum (LGM, 0.020-0.014 Mya; [93,94]). Valdivia interglacial was characterized by the presence of North Patagonian forests and Valdivian rainforests [95], which are habitats associated to *Eupsophus* species [82]. These suitable Late Pleistocene habitats for *Eupsophus* species probably were contracted during periods of glacial advance, whereas distributional range shifted during glacial retreats and warming. Therefore, it is possible hypothesize a wide distribution of *Eupsophus* species during the interglacial, followed by restricted distribution in refugia during the LMG. These cycling events has been hypothesized in other terrestrial vertebrate species [96–98]. Thus, the effect of late Pleistocene cycling events could be related with the actual restricted distribution of some *Eupsophus* species (e. g. *E. migueli, E. altor, E. contulmoensis, E. nahuelbutensis; Eupsophus* sp. *E. septentrionalis*).

Finally, the lineage represented by Villarrica specimens (*Eupsophus* sp.) diverged from *E. roseus* at ∼ 0.088 Mya (Fig 5). Under this temporal scenario it is possible that this lineage lived during interglacial and subsequently was affected by LGM. A central east colonization of an ancestral *E. roseus* population could have given rise to *Eupsophus* sp. during warmer interglacial conditions. In this sense, this putative species probably represents a remnant lineage left behind in central-west Chilean refugia present during LGM. In short, isolation during LGM, the monophyly, and coalescent species delimitation suggest taxonomic differentiation of Villarrica specimens.

Using new molecular datasets and coalescent analyses, our approach revitalizes in an independent way, the hypothesis that *E. roseus* group is composed by eight species. Moreover, we suggest the taxonomic differentiation for Villarica specimens. We suggest filling bioacoustic, morphological, behavioral, and karyotypic data gaps to a deep *Eupsophus* revision.

## Acknowledgements

We thank to Engr. Nicolás González for field assistance. The Corporación Nacional Forestal, Ministerio de Agricultura, Gobierno de Chile allows to collect buccal swabs samples of *Eupsophus* species from wild protected areas (Permit No. 11/2016.-CPP/ MDM/jcr/29.02.2016). Fondecyt 3160328 to EYS-V supported this research.

---

**S1 Table. Sampling locations of *Eupsophus* species**. Coordinates, sample size (N), corresponding species according to Frost [33] and map number from Fig 1 are indicated. Species used as outgroup are also listed (gray cells).

**S2 Table. Sites characterization, partitioning schemes, and nucleotide substitution models for sequences used in this study**. Conservative (C), variable (V), informative (I) and total sites for each marker are indicated. Partitioning schemes, and nucleotide substitution models were determined using Partitionfinder, version 2.1.1 [52].

**S3 Table. Taxonomic index of congruence (C*tax*) calculated for each pair of approaches.** Mean of all the C*tax* values obtained involving a given approach (Mean C*tax*) and total number of species supported by each approach (sp.) is indicated. Species delimitation approaches: Bayesian General Mixed Yule Coalescent model (bGMYC), multi-rate Poisson Tree Processes (mPTP), Tree Estimation using Maximum likelihood, (STEM), Bayesian Species Delimitation (BPP), Multi-locus Species Delimitation using a Trinomial Distribution Model (Tr2), and Bayes factor delimitation (BFD).

**S4 Table. Likelihood scores and Akaike’s information criterion (AIC) results for STEM analysis (see Carstens and Dewey [16]).** Species delimitation scenarios for *Eupsophus* species are indicated in Fig 3A. Species number (sp.), Log-likelihood of the species tree (-lnL), number of parameters (k), AIC, AIC difference (*Δi*), relative likelihood of model given the data (*L*), and the model probabilities (*wi*) are indicated. Note the proximity between –lnL from scenario 11 and 12.

**S5 Table. Bayes factor delimitation results.** Marginal likelihood (MLE) and Bayes factor estimates for species delimitation scenarios indicated in Fig 3A. Species number (sp.) and values using path (PS) and stepping-stone (SS) sampling are indicated

**S1 File. ABGD analyses using COI and concatenated dataset.** Distributions of pairwise distance, ABGD partition, and specimens grouping obtained from COI and concatenated data set are showed.

